## Supplementary Information for "Trait anxiety is associated with hidden state inference during aversive reversal learning"

### Supplementary Methods
#### Differences between studies

**Supplementary Table I:** *Overview of meta-parameters for experiments 1 - 3*

| All collected data sets | | | |
|---|---|---|---|
| | Experiment 1 (fMRI) | Experiment 2 (placebo group from a drug study) | Experiment 3 (new) |
| Visits | 2 | 3 | 1 |
| Sessions per visit | 1 | 1 | 3 |
| Trials per session/visit (bold include included in the joint analysis) | Screening session + training 40 - 60 trials; **Testing session: ~250 trials** | Visit 1: 170 trials **Visit 2: 310 trials** Visit 3: 170 trials | **210 per session (~630 per visit)** |
| Reversal cue trials included (no. trials median, 5th and 95th percentile) | 99 [78 102] | 103 [99 125] | 60/40: 87 [81 110] 75/25: 107 [86 115] 90/10: 100 [92 109] |
| Reversal frequency (trials per phase across all cues) | 30 +/- 2 | 30 +/- 5 | 35 +/- 10 |
| Number of phases | 8-9 | Visit 1: 5-6; Visit 2: 10-11; Visit 3: 5-6 | 6 per session, 18 in total |
| Cues | 3 | 3/visit | 3/session |
| Cue frequency of occurrence (stable-high, stable-low, reversal) | 30-30-40 | 25-25-50 | 20-20-60 |
| Shock probabilities | 25-75% | 25-75% | Session A: 40-60%, Session B: 25-75%; Session C: 10-90% |
| Inter-trial interval (ITI) | 3-5s | 2s | 1-2s |
| Inter-stimulus interval (ISI) | 2-4s | 1s | 1-2s |

**Trait anxiety mean scores**

| | |
|---|---|
| **Study 1** | **40.3** |
| **Study 2** | **37.7** |
| **Study 3** | **41.4** |

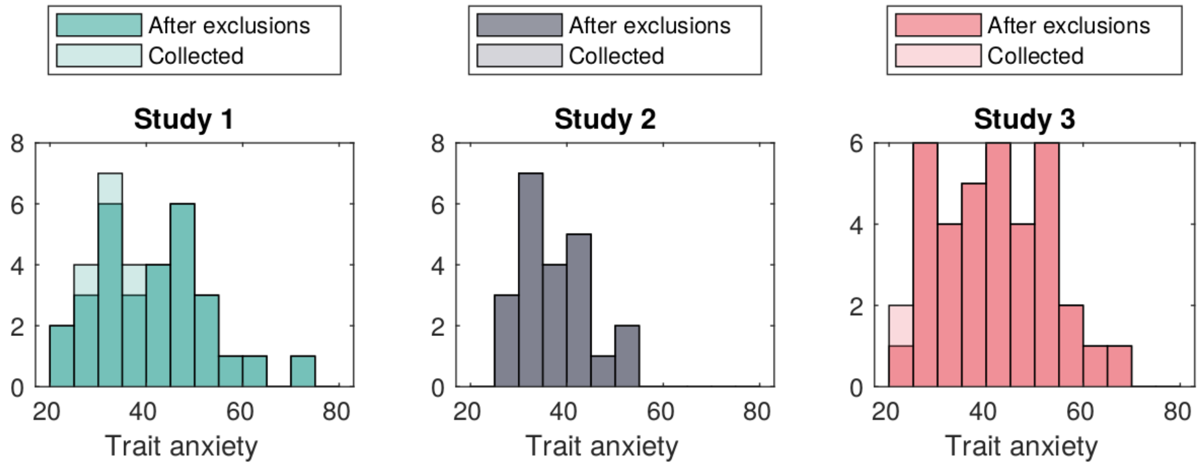


**Supplementary Figure 1** Trait anxiety distributions across the three studies.

#### Inclusion and exclusion criteria

##### *Experiment 1*

*Inclusion criteria*

- Participant is willing and able to give informed consent for participation in the study
- Healthy adults
- Male or Female
- Aged 18 to 40 years
- Not currently taking any medications (except the contraceptive pill)
- Right handed

*Exclusion criteria (if any apply)*

- History of neurological or psychiatric illness (including chronic or remittent pain)
- Contraindications to MRI scanning (including but not limited to a history of claustrophobia, certain metallic implants and metallic injury to eye)
- Pregnancy or are likely to become pregnant during the study
- Recent or current use of psychoactive substances (in the past 3 months)

***Experiment 2***

*Inclusion criteria*

- willing and able to provide informed consent
- male or female, aged 18-40
- body mass index (BMI) of 18-30 kg/m2
- fluent English skills
- non- or light-smoker (< 5 cigarettes a day)

*Exclusion criteria*

- Female participant who is pregnant or breast-feeding
- CNS-active medication during the last 6 weeks
- Current blood pressure or other heart medication (especially aliskiren or beta blockers)
- Intravascular fluid depletion
- Past or present DSM-IV axis-I diagnosis
- Alcohol or substance abuse
- First-degree family member with a history of a severe psychiatric disease
- Impaired liver or kidney function
- Lifetime history of epilepsy or other neurological disease, systemic infection, or clinically significant hepatic, cardiac, obstructive respiratory, renal, cerebrovascular, metabolic, endocrine or pulmonary disease or disorder which, in the opinion of the investigator, may either put the participants at risk because of participation in the study, or may influence the result of the study, or the participant's ability to participate in the study.
- Insufficient English skills
- participated in another study involving certain medication during the last 6 week

***Experiment 3***

*Inclusion criteria*

- Participant is willing and able to give informed consent for participation in the study
- Healthy adults
- Male or Female
- Aged 18 to 40 years
- Not currently taking any medications (except the contraceptive pill).
- Right handed

*Exclusion criteria*

- History of neurological or psychiatric illness (including chronic or remittent pain)
- Pregnancy or are likely to become pregnant during the study
- Recent or current use of psychoactive substances (in the past 3 months)

**Questionnaires**

***Experiment 1***

State-Trait Anxiety Inventory (STAI) [1]

Anxiety Sensitivity Scale (ASI) [2]

Fear of Pain Questionnaire (FPQ-III) [3]

Pain Vigilance and Awareness Questionnaire (PVAQ) [4]

Pain Catastrophizing Scale (PCS) [5]
Centre for Epidemiological Studies Depression Symptoms Index (CES-D) [6]

***Experiment 2***

Eysenck Personality Inventory (EPI) [7]
Beck Depression Inventory (BDI-II) [8]
Trait form of the State-Trait Anxiety Inventory (STAI) [1]
Attentional Control Scale ACS (ACS) [9]
Anxiety Sensitivity Index Revised (ASI-R) [10]
Emotion Regulation Questionnaire (ERQ) [11]

***Experiment 3***

State-Trait Anxiety Inventory (STAI) [1]
Anxiety Sensitivity Scale (ASI) [2]
Fear of Pain Questionnaire (FPQ-III) [3]
Pain Vigilance and Awareness Questionnaire (PVAQ) [4]
Pain Catastrophizing Scale (PCS) [5]
Centre for Epidemiological Studies Depression Symptoms Index (CES-D) [6] .

**Instructions**

While the exact wording of the instructions is slightly different for the three studies, the following key points are maintained:

1. Each of the images is associated with a probability to receive an electrical stimulation
2. This probability can change for any image at any point, so you need to keep paying attention.
3. The breaks in the task don't signal change in probability or restart the task - it is one learning experience all through (*This is slightly different in Study III where we explicitly say that each of the three sessions is starting from a scratch*)

#### Supplementary Note 1: Ratings

***Average ratings in stable cues***

**Supplementary Table 2:** Ratings across stable cues and median-split trait anxiety

| Cue | | 60/40 | | 75/25 | | 90/10 | |
|---|---|---|---|---|---|---|---|
| High-prob | High Anxiety | 60.8% | 61.3% | 72.4% | 73.0% | 82.2% | 85.9% |
| | Low Anxiety | | 60.6% | | 71.7% | | 78.2% |
| Low-prob | High Anxiety | 41.7% | 40.4% | 30.6% | 29.0% | 18.4% | 11.6% |
| | Low Anxiety | | 43.5% | | 32.5% | | 25.8% |

***Ratings in reversal cue***

**Supplementary Table 3:** Ratings across sessions, phases and median-split trait anxiety

| Cue | | 60/40 | | 75/25 | | 90/10 | |
|---|---|---|---|---|---|---|---|
| High-phase | High Anxiety | 63.0% | 63.2% | 71.3% | 71.5% | 76.4% | 77.4% |
| | Low Anxiety | | 62.9% | | 71.2% | | 75.3% |
| Low-phase | High Anxiety | 47.9% | 42.2% | 45.4% | 44.1% | 37.8% | 30.1% |
| | Low Anxiety | | 53.6% | | 46.9% | | 45.5% |

#### Supplementary Note 2: Control and additional analyses

##### Control analysis of slope

To control for the potential effect of pre-reversal baseline on the estimated slopes, we extracted the mean ratings on 5 trials before each reversal and regressed them from the data. We then repeated the analysis reported in the main text focusing on the effect of trait anxiety over the sessions. As reported in Results, we still find the main positive effect of trait anxiety on slopes, $F(1, 146) = 8.59$, $p=.004$, $\eta^2_p=.06$ [.01, .14], as well as interaction between trait anxiety and session, $F(2, 542) = 5.75$, $p=.003$, $\eta^2_p=.02$ [.00, .05], driven by positive association between slopes and TA in the 90/10 condition, slope-beta=7.38, CI95=[3.84 10.91] (all $x10^{-4}$). The association of slope and TA in 90/10 was significant compared to 60/40, $t(460)=-2.39$, $p=.045$ $\eta^2_p=.01$ [.00, .04], and 75/25, $t(574)=-3.31$, $p=.003$ $\eta^2_p=.02$ [.00, .05].

##### Steepness and switch point of the reversal cue

###### *Switch point and steepness*

To isolate subject-specific learning metrics, we used the trial-by-trial expectancy ratings to estimate switch point and switch steepness for each individual change in probability. Only trials from the reversal cue were included in this analysis. For each switch, we extracted 5 trials prior and 10 trials post reversal and demeaned the time-series. Using this 'chunk', we determined the point of steepest change ('switch point') by calculating the cumulative sum and identifying the point of highest (high-to-low switch) or lowest (low-to-high switch) value in the series.'' Next, a smaller chunk of 10 trials (5 preceding and 5 following) around the identified switch point was extracted. A sigmoidal curve (Eq. 1) with a free parameter for steepness was fitted to this smaller chunk. ***X*** corresponds to the time series of 10 trials, ***b*** is the inflection point which was fixed to ***b***=*5.5* (midpoint) and ***a*** represents the steepness.

$$\text{Eq. 1}\ f(x, a, b) = \frac{1}{1 + e^{-a(x-b)}}$$

The fitted value for ***a*** was recorded and log transformed before being used as an estimate for switch steepness in the main analysis.

As reported in the main text, the reversal time of the switching cue was semi-random and not cued, and participants had to infer reversals from observations. This was likely to cause delays before updating could begin, and to vary between participants and even between reversals. Because averaging over variable time courses can lead to incorrect conclusions [e.g., 12], we estimated the time-point of the steepest change post-reversal separately for each participant and reversal. This also allowed us to estimate the steepness at that point using a sigmoidal fit. This analysis provided an estimate of how many trials after the reversal ratings began to change as well as an estimate of how fast ratings changed once the change began (steepness). Splitting data by estimated switch point and steepness (Supp Figs 2a and 2b) showed that our approach indeed captured variability in these two aspects.

Analyzing the steepness of estimated switches, it increased across sessions (-2.30, -2.10 and -1.4 on log scale), as reflected in a main effect of session, $F(2, 205)=20.80$, $p<.001$, $\eta^2_p=.16$ [.09, .25]. This was driven by significantly higher steepness in the 90/10 condition compared to 60/40, $t(226)=-6.14$, $p<.001$ $\eta^2_p=.14$ [.06, .22], and to 75/25, $t(181)=-4.61$, $p<.001$, $\eta^2_p=.10$ [.04, .19]. The steepness was also positively associated with trait anxiety TA, $F(1,94)=7.39$, $p=.008$ $\eta^2_p=.07$ [.01, .19], indicating that more anxious participants adjusted their shock probability ratings faster than less anxious participants. Turning to analysis of the estimated switch point. The average switch occurred 2.91, 2.90 and 3.28 trials after the true reversal (60/40, 75/25, 90/10 respectively). There was no relationship between the estimated switch point, session and TA. This indicates that while high trait anxious participants performed more abrupt switches, these did not occur earlier or later compared to individuals lower in trait anxiety, and this was true for all sessions.

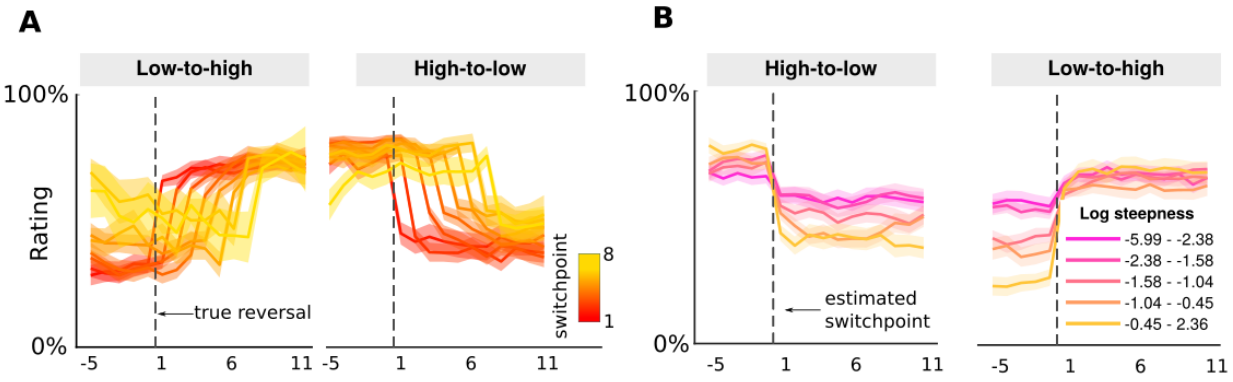


**Supplementary Figure 2 Time courses of different values of switchpoint and steepness.** The time courses sorted by (**a**) estimated switch point, locked to the true reversal, and (**b**) estimated switch steepness, locked to the estimated switch point. Thick lines in all panels show mean across the N=89 participants. Shaded areas show standard error of the mean.

#### Relationship between stable cues and the reversal cue

To check whether the anxiety effects in stable cues and the two phases of the reversal cue were related, we correlated the difference in stable-high and stable-low cues (“stables difference”) with the difference in low- and high phases (“phase difference”). There was a positive association in all three sessions ($r(35)_{60/40}=0.20$, CI=[-.01,.39], $p=.066$; $r(87)_{75/25}=0.24$, CI=[-.10,.52], $p=.165$; $r(36)_{90/10}=0.49$, CI=[.20,.70], $p=.002$), but only the correlation in the 90/10 condition was significant after correction.

#### Meaningful versus oddball learning

To assess the impact of the selection window more systematically, we used alternative decision criteria (i.e., cut-offs at trials 5. 7, 10 and 13) to generate meaningful/oddball trials (see the calculated learning rates for the four analyses plotted separately for each condition below). Notably, the different cutoffs produced very similar results. They all clearly capture the main effect difference between meaningful and oddball learning. Indeed, using a formal LMM, there was no significant main effect or interaction involving the cutoff, all ps > 0.9.

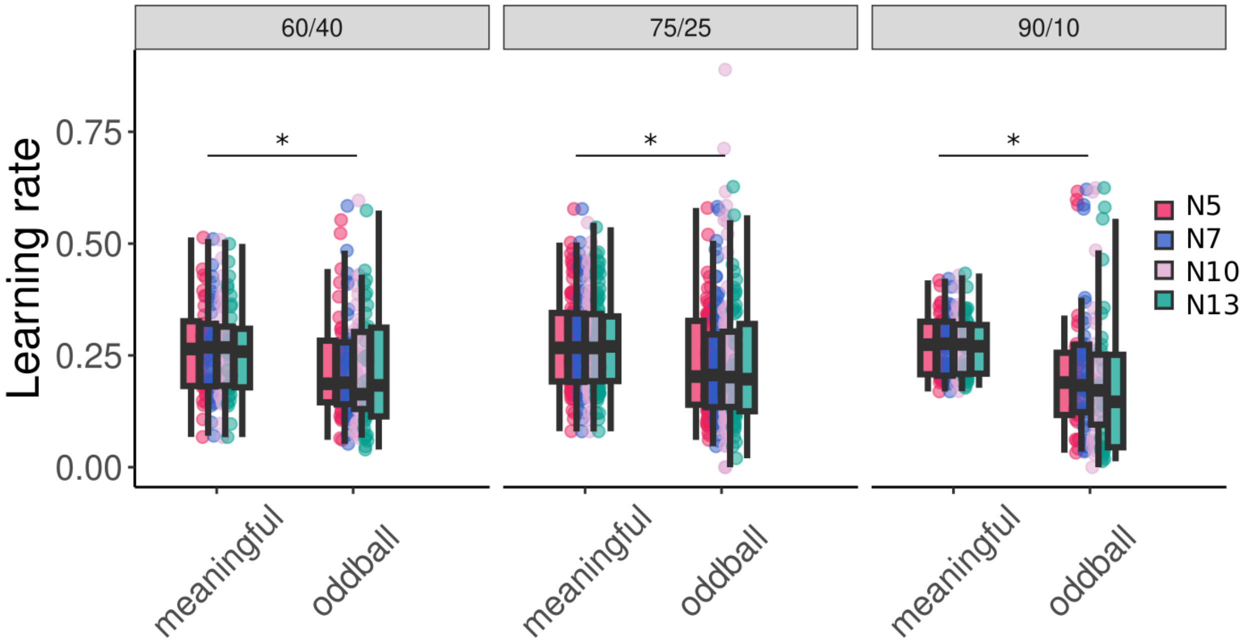


**Supplementary Figure 3 Selection window for oddball categorization.** Model-free learning rates categorized by oddball (surprising events during relatively stable periods that don't signal change in environment) and meaningful trials (trials following a true reversal)**.** The different colors represent different cutoff points used to classify meaningful and oddball trials (5-7-10-13). The three conditions contain N=36, N=88 and N=37 participants. Boxplots indicate the median and interquartile range (IQR, i.e., distance between the first and third quartiles). The lower and upper hinges correspond to the first and third quartiles (the 25th and 75th percentiles). The upper whisker extends from the hinge to the largest value no further than 1.5* IQR from the hinge. The lower whisker extends from the hinge to the smallest value at most 1.5* IQR of the hinge.

##### Internal parameters of the winning model

To better understand the fitted n-state model, we explored it's behavior and fitted parameters in more detail. First, we analyzed whether the number of states the fitted model created was related to TA, but this was not the case (mean number of states high TA: 1.82 vs. 1.81 in low TA). Second, we investigated the timepoint when the model tended to switch states. A corresponding LMM found no effect of session or anxiety, consistent with our behavioral results reported above (see "estimated switch point"; model inferred switch points were 2.93, 3.20 and 3.19 trials after the true reversal, for the three sessions). Next, we analyzed the fitted step sizes for the positive and negative outcomes $\tau^{+}$and $\tau^{-}$. A LMM with parameter type ($\tau^{+}/\ \tau^{-}$), TA and session as fixed effects found a significant main effect of outcome type, $F(1,228)=37.03$, $p<.001$, $\eta^2_p=.14$ [.07, .22], which reflected that shocks elicited larger updates than no-shocks $\tau^{+}=1.13$ vs. $\tau^{-}=0.73$). There was no main effect of TA, $F(1,83)=.21$, $p>.05$, $\eta^2_p=.00$ [.00, .06], or interaction of outcome type and TA, $F(1, 228)=.16$, $p>.05$, $\eta^2_p=.00$ [.00, .02]. Note that the same two parameters of the 1-state model, had a similar difference $\tau^{+}$=1.17 vs.$\tau^{-}$=0.86), suggesting that differential learning from shock and no-shock events alone was unable to explain our behavioral effects of TA. Lastly, no effects of session, TA or interaction where found when analyzing the remaining parameters $\eta$ (switch threshold), $\alpha_0$ and $\beta_0$.

***Supplementary Table 4a: Estimated parameters for 1-state model***

| | 60/40 | | | 75/25 | | | 90/10 | | |
|---|---|---|---|---|---|---|---|---|---|
| | Stable-high (N = 36) | Stable-low (N = 36) | Reversal (N = 36) | Stable-high (N = 88) | Stable-low (N = 88) | Reversal (N = 88) | Stable-high (N = 37) | Stable-low (N = 37) | Reversal (N = 37) |
| **Tau shock** | 1.05 ± 0.72 | 0.89 ± 0.73 | 0.94 ± 0.72 | 1.17 ± 0.75 | 0.86 ± 0.68 | 1.15 ± 0.70 | 1.43 ± 0.71 | 0.68 ± 0.65 | 1.19 ± 0.74 |
| **Tau noshock** | 0.86 ± 0.72 | 0.99 ± 0.67 | 0.77 ± 0.58 | 0.70 ± 0.62 | 1.15 ± 0.75 | 0.74 ± 0.57 | 0.71 ± 0.81 | 1.49 ± 0.73 | 0.85 ± 0.69 |
| **Alpha0** | 5.06 ± 3.29 | 4.18 ± 3.22 | 5.07 ± 3.52 | 4.95 ± 3.37 | 4.39 ± 3.39 | 5.33 ± 3.24 | 4.69 ± 3.63 | 3.08 ± 2.51 | 5.55 ± 3.31 |
| **Beta0** | 5.03 ± 3.38 | 4.54 ± 3.23 | 4.51 ± 3.27 | 4.49 ± 3.10 | 5.14 ± 3.33 | 4.90 ± 3.30 | 3.37 ± 3.04 | 4.29 ± 2.98 | 3.83 ± 3.01 |
| **Lambda** | 0.88 ± 0.16 | 0.93 ± 0.09 | 0.87 ± 0.16 | 0.88 ± 0.17 | 0.89 ± 0.13 | 0.87 ± 0.11 | 0.89 ± 0.22 | 0.94 ± 0.07 | 0.85 ± 0.13 |

***Supplementary Table 4b: Estimated parameters for n-state model***

| | 60/40 | | | 75/25 | | | 90/10 | | |
|---|---|---|---|---|---|---|---|---|---|
| | Stable-high (N = 36) | Stable-low (N = 36) | Reversal (N = 36) | Stable-high (N = 88) | Stable-low (N = 88) | Reversal (N = 88) | Stable-high (N = 37) | Stable-low (N = 37) | Reversal (N = 37) |
| **Tau shock** | 1.26 ± 0.75 | 0.99 ± 0.67 | 0.98 ± 0.64 | 1.18 ± 0.70 | 0.85 ± 0.67 | 1.11 ± 0.70 | 1.42 ± 0.74 | 0.61 ± 0.73 | 1.16 ± 0.74 |
| **Tau noshock** | 0.88 ± 0.73 | 1.13 ± 0.65 | 0.77 ± 0.57 | 0.66 ± 0.67 | 1.16 ± 0.68 | 0.63 ± 0.54 | 0.65 ± 0.76 | 1.40 ± 0.72 | 0.66 ± 0.51 |
| **Alpha0** | 5.90 ± 3.20 | 5.21 ± 3.26 | 5.73 ± 3.33 | 5.13 ± 3.05 | 5.35 ± 3.37 | 4.86 ± 3.30 | 5.91 ± 3.12 | 4.78 ± 3.31 | 4.90 ± 3.26 |
| **Beta0** | 5.94 ± 2.97 | 5.58 ± 3.21 | 5.07 ± 3.28 | 5.40 ± 3.05 | 5.36 ± 3.08 | 4.44 ± 2.90 | 5.67 ± 3.38 | 5.01 ± 3.12 | 4.42 ± 3.43 |
| **Lambda** | 0.85 ± 0.25 | 0.94 ± 0.07 | 0.89 ± 0.17 | 0.91 ± 0.18 | 0.91 ± 0.15 | 0.90 ± 0.10 | 0.90 ± 0.25 | 0.97 ± 0.04 | 0.90 ± 0.10 |
| **Eta** | 4.92 ± 3.13 | 4.80 ± 3.09 | 4.67 ± 2.64 | 4.34 ± 2.96 | 4.12 ± 2.82 | 5.38 ± 2.73 | 3.76 ± 3.20 | 4.65 ± 3.41 | 4.45 ± 2.54 |

#### Experienced uncertainty

One possibility is that a drive to reduce uncertainty in high TA motivates increased state inference. We used the standard deviation of the current state (calculated based on trial-specific alpha and beta) summed across trials as an overall estimate of uncertainty. Using a LMM, we then explored the relationship between uncertainty and anxiety. The model found no significant association between uncertainty and anxiety, or interaction with session. Albeit not significant, there was a general trend towards a negative association between TA and experienced uncertainty in the 90/10 condition as one would expect from an agent which successfully infers state structure (see plot below). We also found a weak positive association between TA and uncertainty in the 60/40 condition.

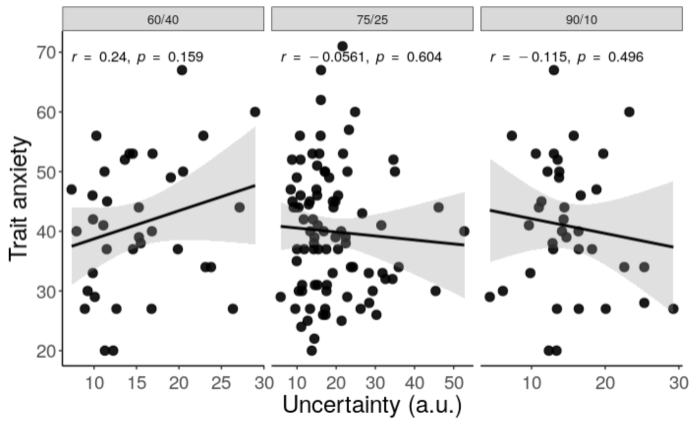


**Supplementary Figure 4 Experienced uncertainty.** Relationship between experienced uncertainty (estimated by the n-state model) and trait anxiety separately for the three sessions. Trait anxiety did not relate to experienced uncertainty in any of the sessions. The three panels include N=36, N=88 and N=37 data points. Shown correlation coefficients reflect Pearson's correlation (two-tailed) and uncorrected p-values.

#### Differences between studies

To compare difference between studies we ran a number of analyses using the 75/25 condition. We did not find any significant difference between studies. However, the trait anxiety scores were generally lowest in the drug study. Therefore, there are only 6 high TA participants in Study 2. The uncertainty in high TA – low phase condition is too high to conclude whether we see the same effect as in the other two studies. Apart from the behavioral data, we also looked at differences between studies in the slope after reversal. As reported in the main text we see a main positive effect of trait anxiety. There is no interaction between study and TA.

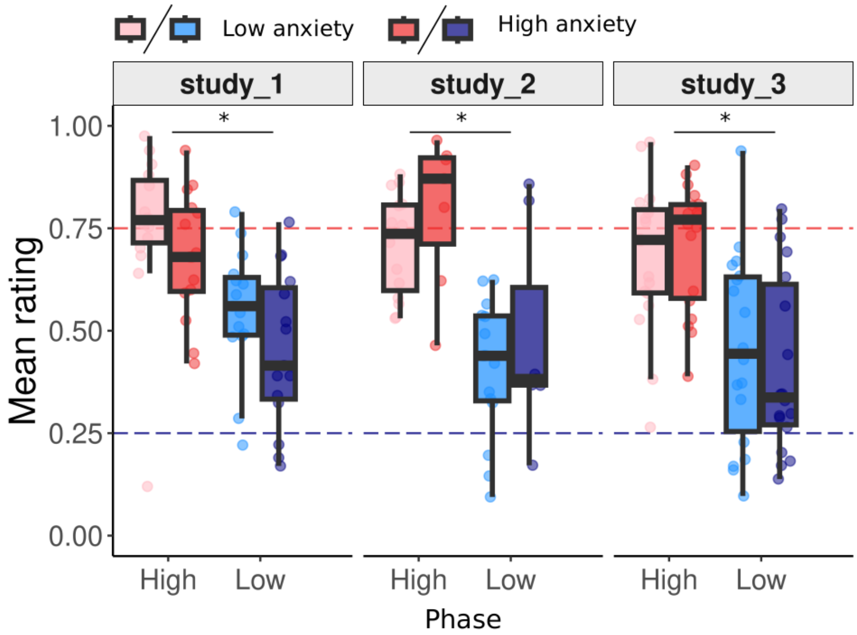


**Supplementary Figure 5 Ratings across studies in the 75/25 session.** Mean ratings for trials 10+ after reversal split by phase (high/low), session (60/40, 75/25, 90/10) and median-split trait anxiety. Statistical test was performed using LMM model and follow-up ANOVAs. The three conditions contain N=36, N=88 and N=37 participants. Boxplots indicate the median and interquartile range (IQR, i.e., distance between the first and third quartiles). The lower and upper hinges correspond to the first and third quartiles (the 25th and 75th percentiles). The upper whisker extends from the hinge to the largest value no further than 1.5* IQR from the hinge. The lower whisker extends from the hinge to the smallest value at most 1.5* IQR of the hinge.

#### Modeling

##### *Percentage of participants best fitted by each model*

| | 60/40 | 75/25 | 90/10 |
|---|---|---|---|
| 1-state | 41.7% | 43.2% | 37.8% |
| n-state | 58.3% | 56.8% | 62.2% |

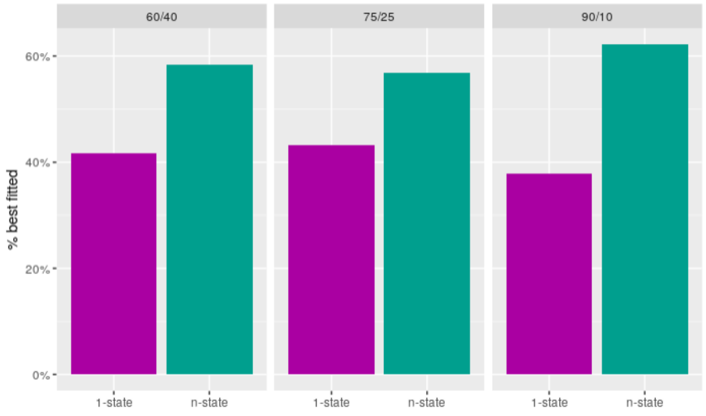


**Supplementary Figure 6 Proportions of participants best fitted by each model.** Percentage of participants best fitted by the 1-state (purple) and the n-state (green) models separately for each session.

##### *Best model fit by condition*

To assess how model fit developed within individual over the three sessions we binned participants according to the best fitting model in each condition. In the plot below "1" stands for gradual (1-state) learning while "n" stands for structure learning (n-state). For example, "1-1-n" therefore represents a case where in the 60/40 and 75/25 conditions the 1-state model fitted best and in the 90/10 condition the n-state model fitted best. Summarizing data in such a way allows us to assess strategy switches between sessions. We plot the results below.

Interestingly, there appears to be a high degree of internal consistency - bins with the same strategy across the three sessions (1-1-1, n-n-n) seem to stand out (38%; chance level 11.1%). This is followed by a group of participants which relied on gradual learning in the noisier conditions but that employed structure learning in the 90/10 condition (1_n_n and 1_1_n; together 32%).

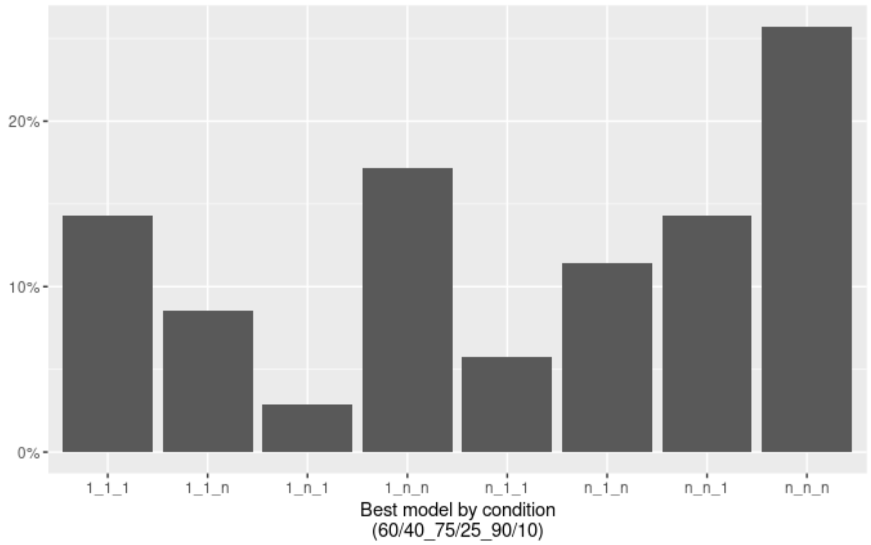


**Supplementary Figure 7 Distribution of individuals according to which model fitted best across the three sessions.** All x axis labels occur in the 60/40-75/25-90/10 order.

Next, we correlated the relative model fit for participants who completed all three sessions (correlation matrix below). This analysis further confirmed within-participant consistency of relative n-/1-state model fit, particularly between 75/25 and 90/10 (r=.44, p=.007), and 60/40 and 75/25 (r=.38, p=.023), but not between 60/40 and 90/10

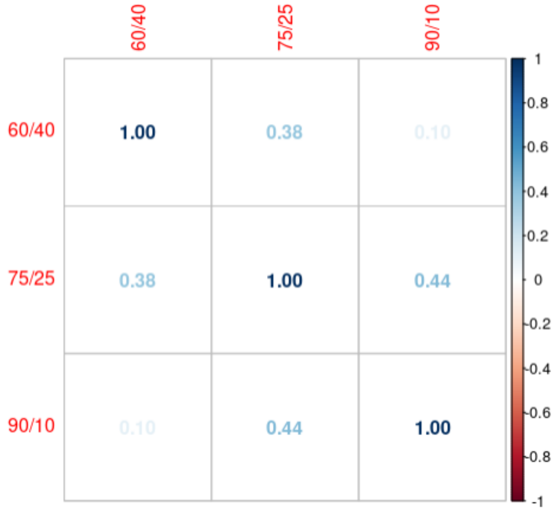

.

**Supplementary Figure 8 cross-session correlations of model fits.** Correlation ofn relative model fits (difference of BIC scores for the n-state and 1-state models) across the three sessions. The displayed numbers represent Pearson's correlation coefficients.

##### *Out-of-sample model fit and behavioral markers*

In the main text we report the relationship between two key markers of state inference (slope and differential learning from oddballs compared to trials after reversal). To check generalizability of model fit in relation to these behavioral markers we fitted the models to the first half of trials (phases 1-3) and correlated the relative model fit with behavioral measures derived from second half of each session (phases 4+). In both cases, the behavioral marker from the second half correlated with the first half relative model fit. Specifically, the Spearman's rho correlation coefficients were r=.332, p=.002 for slopes and r=.21, p=.046 for meaningful/oddball.

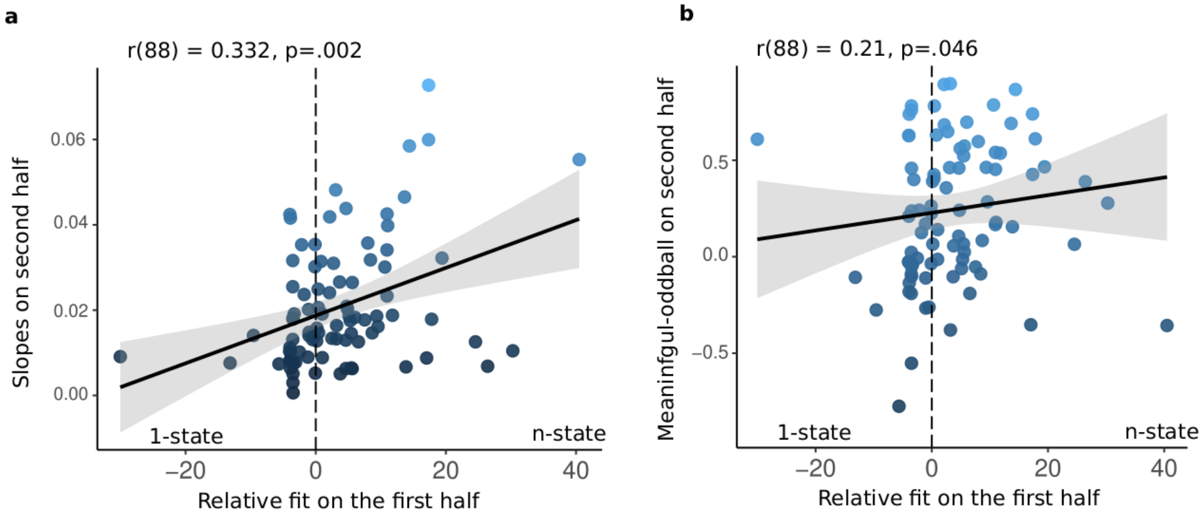


**Supplementary Figure 9 Out of sample correlations.** Correlation between relative model fit (BIC difference between n-state and 1-state models) from the first half and (**a**) estimated slopes, r(87)=.33, p=.002, CI=[.13, .51] and (**b**) the difference between meaningful and oddball trials, r(87)=.21, p=.046, CI=[.01, .41], both from the second half. Both analyses of N=89 participants were performed using Spearman correlation, two-sided. Displayed line and shading corresponds to linear model fit, y ~ x.

##### *Order effects*

We investigated whether the order of sessions impacted the relative fit of gradual and state inference model.. Specifically, focusing on order effects in the relationship between model fit and trait anxiety, we found no significant main effect or interaction of order. We present the sessions by the order they occurred in in the figure below. Additionally, we specifically investigated the tendency to use state inference in the 60/40 condition depending whether this occurred first or whether it was preceded by 90/10. The mean reliance on state inference was numerically higher when 90/10 preceded 60/40 (deltaBIC = 16.4) compared to when 60/40 occurred first (deltaBIC = 6.86), but this effect was not statistically significant, t(42)=1.61, p=.11, $\eta^2_p$=.06 [.00, .23] (Supp Fig 10a).

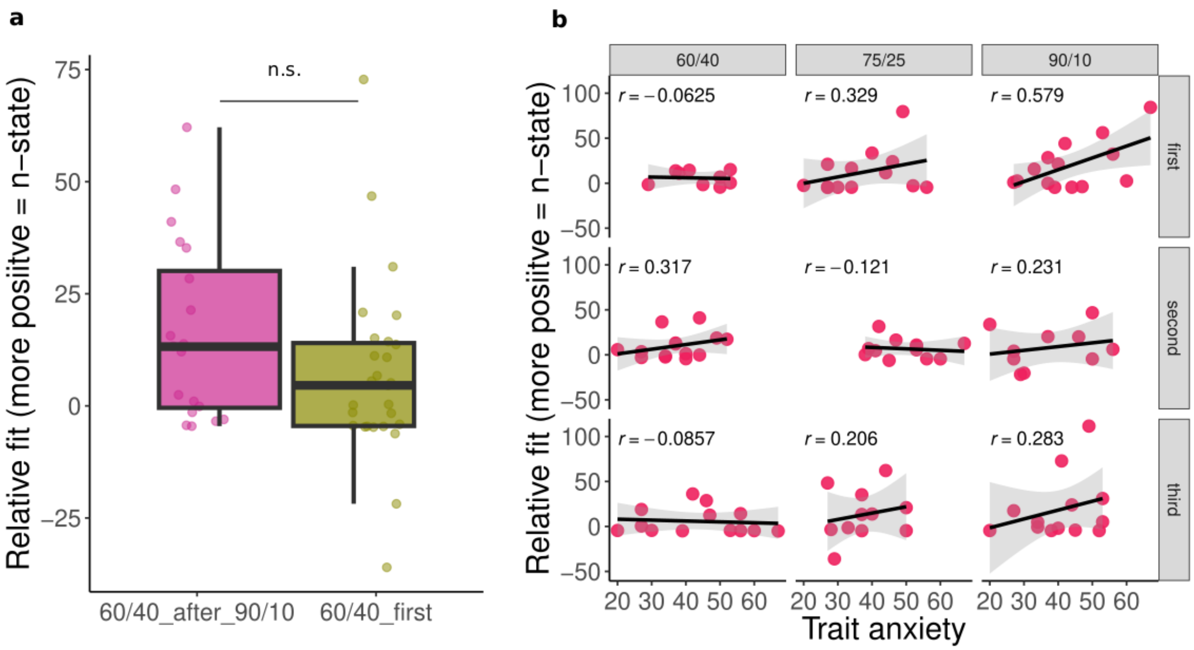


**Supplementary Figure 10 Impact of session order on relative fit and relationship with trait anxiety.** (**a**) Model fit in the 60/40 session tends to (not significantly) favor state inference (positive values) if the session was preceded by the 90/10 session, t(42)=1.61, p=.11, $\eta^2_p$=.06 [.00, .23], (N=20) compared to when it occurred first (N=26). (**b**) Correlations between trait anxiety and relative model fit split by session and in which order it occurred, assessed by Pearson correlation. The plots include the following number of data points: In 60/40 N=9, N=14, N=13; in 75/25 N=13, N=12, N=11; in 90/10 N=14, N=10, N=13 (all in the logical order: first, second, third). Displayed line and shading corresponds to linear model fit, y ~ x. Boxplots in panel a) indicate the median and interquartile range (IQR, i.e., distance between the first and third quartiles). The lower and upper hinges correspond to the first and third quartiles (the 25th and 75th percentiles). The upper whisker extends from the hinge to the largest value no further than 1.5* IQR from the hinge. The lower whisker extends from the hinge to the smallest value at most 1.5* IQR of the hinge.

#### Raw data for most gradual and SI participants

To verify that the two models capture gradual (1-state model) and state inference (n-state) behavior we calculated the BIC score difference between 1- and n-state models for each participant. As reported in the main text (Fig. 6a) participants better fitted by the n-state showed steeper learning. Here, we extend this report by presenting the raw data of the "most gradual" and "most switchy" participants (See Supp. Fig 11). Visual inspection of the data helps us further validate the model fits: participants with positive BIC difference (green) show large jumps in contingencies following reversal while participants with negative BIC difference show shallower learning curves.

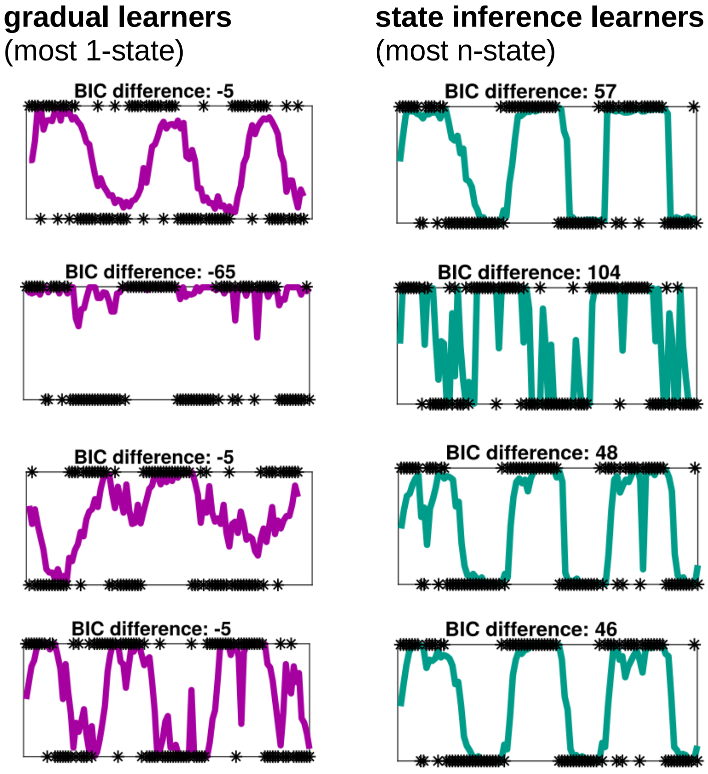


**Supplementary Figure 11 most gradual and state-learning participants.** Raw time-courses of participants best fitted by the gradual learning 1-state model (left) versus participants best fitted by the state switching model (n-state; right)

#### 1-state model, Pearce-Hall and Rescorla-Wagner

The 1-state model behaves very similarly to commonly used gradual learning models such as Pearce-Hall [13,14]. We opted to use the beta-based model for gradual learning as this allowed us to minimize technical differences between the 1-state and n-state models. To check that the 1-state model performs similarly to Pearce-Hall (PH) and Rescorla-Wagner (RW; later 'associative learning models') we fitted both models to the data. Because the 1-state model and the associative learning models use different likelihood functions (the beta-based models use the beta distribution likelihood while the associative learning models use the root mean squared error) we used likelihood-independent measure to assess model fit: correlation between data and model predictions. Using this measure we found no difference between models, $F(2,379)=2.04$, n.s. (Supp. Fig. 13a). Next, we used the fitted parameters to generate posterior predictive checks for all three models (Supp. Fig. 13b). These show that all three models in fact fit the data in a very similar manner.

An interesting aspect of the 1-state model is that it can closely approximate both RW and PH models. The main difference between the associative learning models is stationarity of the learning rate. In RW, the learning rate is the same for all trials while in PH it changes depending on the previous prediction error (i.e., 'associability'). Under PH, learning rates increase after reversal and decay as prediction errors decrease over time. In the case of our beta-based 1-state model, a constant amount $\tau$ is added either to alpha or beta on each trial. While intuition might suggest that this results in constant learning rates, in fact, learning rates behave very similarly to those of PH - larger updates occur after reversal. For demonstration of this, see Supplementary Figure 12.

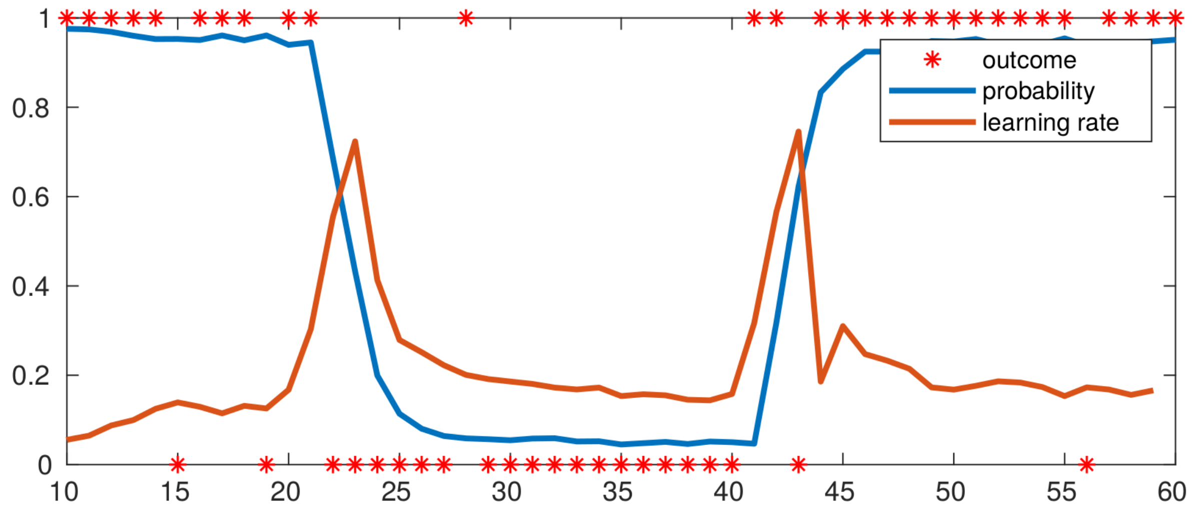


**Supplementary Figure 12 learning rates and probability estimates simulated using the 1-state model.** Data generated using the 1-state beta model (blue) with parameters $\alpha_0 = 4$, $\beta_0 = 4$, $\lambda = 0.8$, and $\tau^{+/-} = 0.5$. Learning rates calculated for each trial show an initial bump after contingency reversal followed by a decay (red). Averaged over 500 simulations.

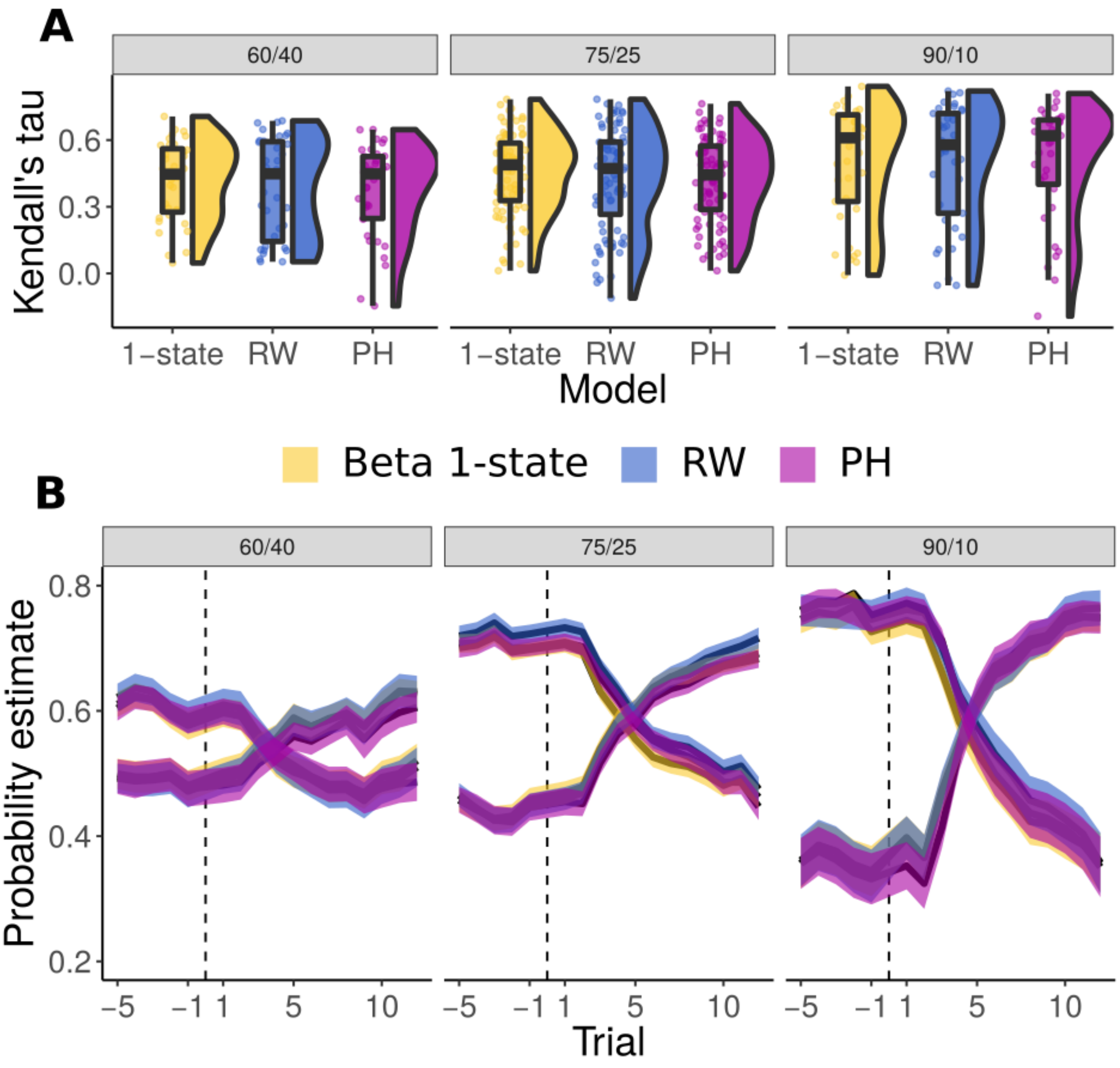


**Supplementary Figure 13 Beta-based models versus reinforcement learning models.** Comparison of Rescorla-Wagner (blue) and Pearce-Hall (purple) models as examples of associative learning models with the 1-state beta learner (yellow). (**a**) Distribution of Kendall correlation coefficients between model prediction and raw data (one for each participant). The data were analyzed using LMM which found a main effect of session, $F(2,401)=22.77$, $p<.001$, , $\eta^2_p=.10$ [.05, .16], but no difference or interaction of a model. (**b**) Posterior predictive checks show that all three models capture behavior in a similar manner. Boxplots in panel a) indicate the median and interquartile range (IQR, i.e., distance between the first and third quartiles). The lower and upper hinges correspond to the first and third quartiles (the 25th and 75th percentiles). The upper whisker extends from the hinge to the largest value no further than 1.5* IQR from the hinge. The lower whisker extends from the hinge to the smallest value at most 1.5* IQR of the hinge. In panel b) the thick lines indicate mean while shaded areas indicate standard error of the mean. Both panels include N=35, N=88 and N=37 participants (the three sessions respectively).

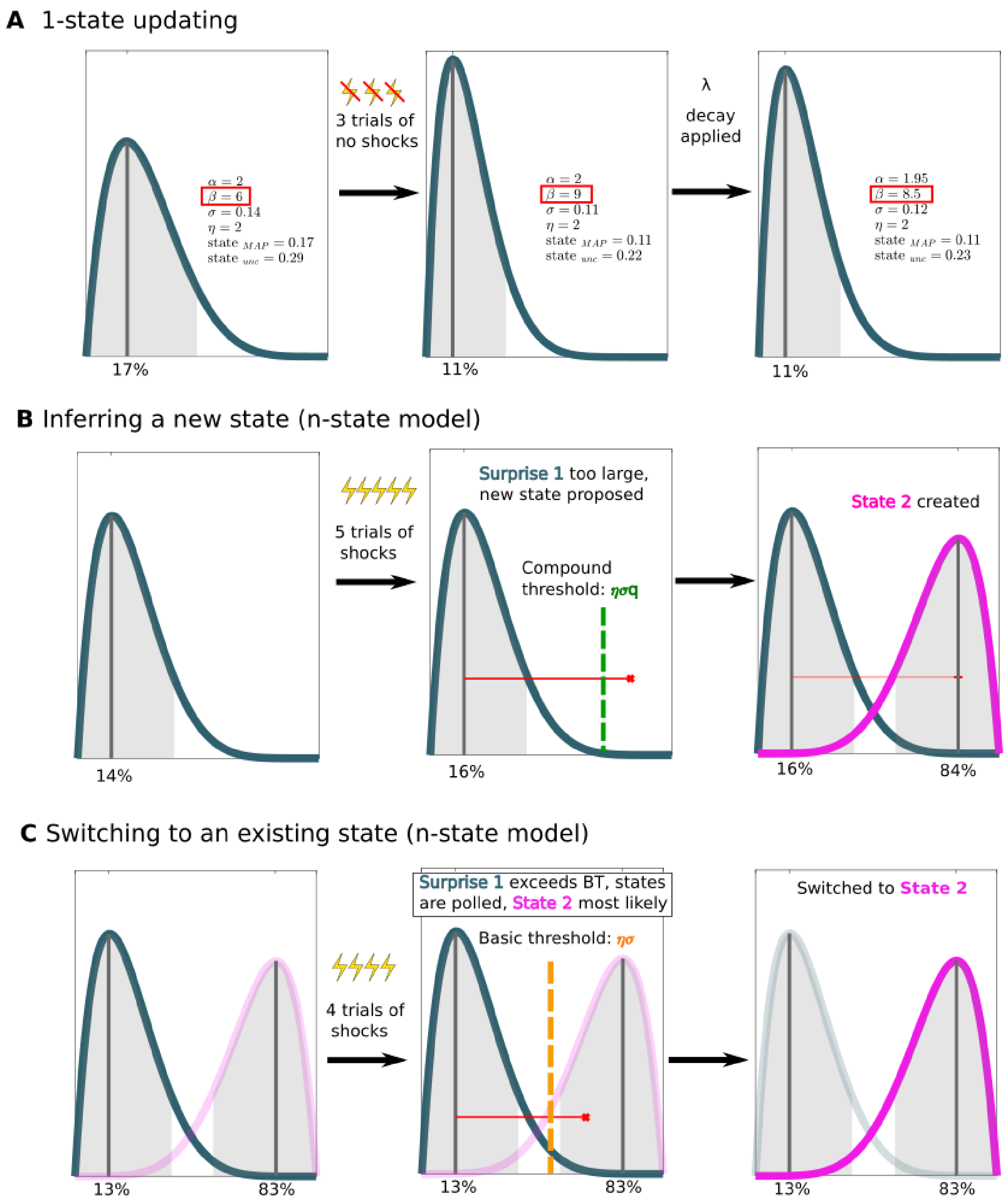


**Supplementary Figure 14 Mechanisms of beta updating.** (**a**) Example of gradual updating of one state where the beta parameter increases after receiving three no-shock outcomes. Consequently, the probability estimate (mode) decreases as does the state uncertainty σ. The right panel shows an example of applying the decay parameter λ - in the actual model this happens on every trial. (**b**) Example of the running cumulative surprise (red line) exceeding the compound threshold. Since there is no existing state, state 2 is created. (**c**) Situation where the model already has two states in memory and the basic threshold for the current State 1 is exceeded. Under this scenario the model polls for the state with less surprise. Focusing on the middle panel, State 2 is currently closer to the proposal value (current state value + surprise) which results in the switch to State 2 (right panel).

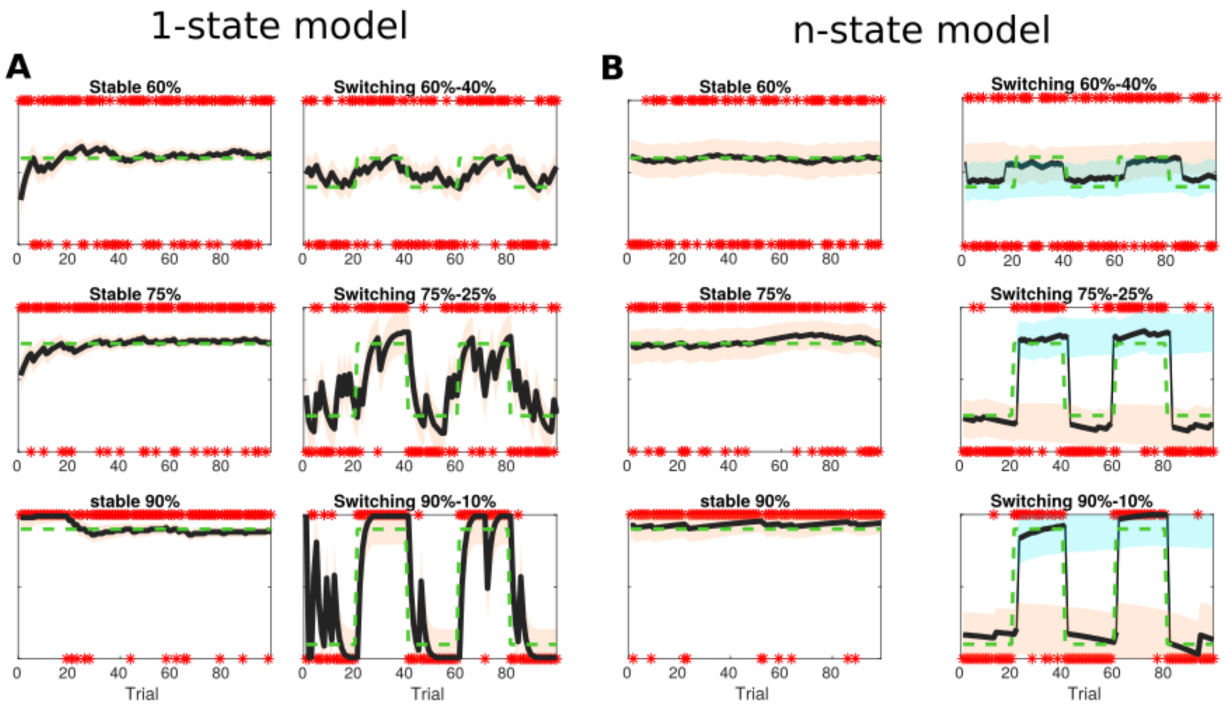


**Supplementary Figure 15 Model fit to synthetic data**. Fit of the 1- (**a**) and n-state (**b**) models to synthetic data. Left panels in each section show fit to stimulus with fixed probability (mirroring the stable cues), right panels show fit to data with changing shock probabilities. Shaded areas represent uncertainty of each state.

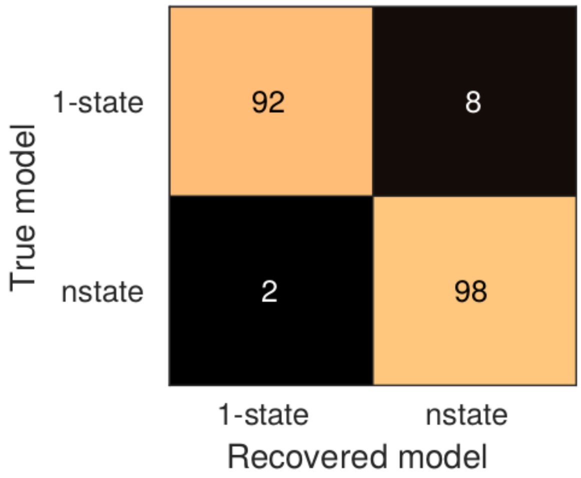


**Supplementary Figure 16.** Confusion matrix of model recovery. Numbers represent sum of datasets (n=100) generated and recovered by a given combination of models.
